# Mechanical Drivers of Glycosaminoglycan Content Changes in Intact and Damaged Human Cartilage

**DOI:** 10.1101/2024.06.17.599262

**Authors:** Seyed Ali Elahi, Rocio Castro-Viñuelas, Petri Tanska, Lauranne Maes, Nele Famaey, Rami K. Korhonen, Ilse Jonkers

**Affiliations:** Department of Movement Sciences, KU Leuven, Leuven, Belgium; Mechanical Engineering Department, KU Leuven, Leuven, Belgium; Department of Development and Regeneration, Skeletal Biology and Engineering Research Center, KU Leuven; Department of Technical Physics, University of Eastern Finland, Kuopio, Finland; University Hospitals Leuven, Division of Rheumatology, Leuven, Belgium

## Abstract

Articular cartilage undergoes significant degeneration during osteoarthritis, currently lacking effective treatments. This study explores mechanical influences on cartilage health using a novel finite element-based mechanoregulatory model, predicting combined degenerative and regenerative responses to mechanical loading. Calibrated and validated through one-week longitudinal ex vivo experiments on intact and damaged cartilage samples, the model underscores the roles of maximum shear strain, fluid velocity, and dissipated energy in driving changes in cartilage glycosaminoglycan (GAG) content. It delineates the distinct regenerative contributions of fluid velocity and dissipated energy, alongside the degenerative contribution of maximum shear strain, to GAG adaptation in both intact and damaged cartilage under physiological mechanical loading. Remarkably, the model predicts increased GAG production even in damaged cartilage, consistent with our in vitro experimental findings. Beyond advancing our understanding of mechanical loading’s role in cartilage homeostasis, our model aligns with contemporary ambitions by exploring the potential of in silico trials to optimize mechanical loading in degenerative joint disease, fostering personalized rehabilitation.

## Introduction

Osteoarthritis (OA), the most common joint disease, affects more than 535 million people worldwide in 2020 (1), causing joint pain and reduced mobility (2). With an ageing population, the burden of OA is expected to increase, causing an annual reduction of 15 million Quality-Adjusted Life Years, comparable to conditions such as cardiovascular disease and cancer (3). As a result, the development of successful curative or preventive treatment strategies for OA remains a major unmet challenge (4). OA is a multifactorial joint disease with associated changes in the articular cartilage, subchondral bone, and synovial inflammation (5). Cartilage degeneration and loss of its complex mechanobiological properties are the hallmarks of OA (6). There is a critical gap in understanding how mechanical micro-environment intricately interact with biological changes in cartilage, hindering the progress in creating effective treatment strategies. This understanding is in particular essential for informing interventions aimed at limiting cartilage degeneration in OA and promoting enhanced cartilage regeneration to restore optimal joint function, the ultimate ambition of personalized rehabilitation or exercise prescription.

Articular cartilage provides structural support, absorbs impact, and ensures smooth articulation (7). The complex, multiphasic (*i*.*e*., solid-fluid mixture) microstructure of the extracellular matrix (ECM) produced by chondrocytes, is critical for weight-bearing functions and chondrocyte integrity (8). It consists of proteoglycans (PGs) and collagen fibers arranged in depth-dependent arcade shapes. The PGs, composed of glycosaminoglycan (GAG) chains covalently linked to a core protein, are highly negatively charged, and form a hydrated gel-like structure that resists compression. The interaction between GAGs, collagen fibers and the interstitial fluid provides a unique mechanical behavior that enables cartilage to withstand loading during locomotion (9) and maintain cartilage homeostasis.

Mechanical loading plays a pivotal role in maintaining the ECM constitution. Physiological mechanical loading assists the chondrocytes in maintaining cartilage integrity through fluid flow and subsequent nutrient delivery and enhances the chondrocyte anabolic activity through energy dissipation (9–12). The depth-dependent ECM constitution has been proposed to adequately modulate the mechanical environment of the chondrocytes throughout the cartilage thickness to maintain their phenotype and local ECM production (13).

However, when cartilage is structurally degraded and/or exposed to abnormal mechanical loading, the tissue is subjected to excessive localized strains and stresses compromising the mechanical microenvironment of the chondrocytes. In addition to the induction of catabolic cellular processes, this leads to ECM disorganization and mechanical failure of the ECM constituents (14), triggering a degenerative cascade in cartilage including collagen fibril degradation and fibrillation, PG depletion, and changes in tissue hydration (15). The associated increase in fluid velocity will cause small protein fragments (e.g. GAGs) to extrude from the cartilage (16, 17) and further disorganize the collagen fibrils, thereby further accelerating ECM degeneration (18–20). To date, it is unknown how the local mechanical micro-environment (e.g. tissue strain or fluid flow) contributes to ECM adaptation under varying mechanical loading or cartilage mechanical condition (e.g. in the presence of a defect, etc.). Elucidation of the predominant factors influencing ECM adaptation to loading in intact and damaged cartilage would facilitate the development of individualized therapeutic approaches to effectively mitigate these ECM changes to prevent cartilage degeneration and even restore cartilage function.

Physics-based modeling techniques are promising approaches to investigate the complex interplay between mechanical loading on the one hand and time- and depth-dependent changes in the cartilage constituents on the other. In particular, adaptive finite element (FE) models allow relating cartilage macro-level loading to local tissue mechanics (strains and stresses), as well as the associated fluid mechanics. In combination with a time-dependent adaptive algorithm, the depth-dependent degeneration of the cartilage ECM can then be predicted based on constituent failure criteria (21). More specifically, the stresses or strains on the ECM constituents are compared to experimentally determined thresholds and when the threshold is exceeded, depth-dependent failure of the ECM constituents is predicted, which changes the local cartilage structure, composition and/or mechanical properties, indicating tissue degeneration. We have recently introduced a Cartilage Adaptive REorientation Degeneration (CARED) model that integrates all experimentally observed mechanics-driven cartilage degeneration mechanisms, more specifically collagen fibril reorientation, collagen loss, PG depletion, and changes in tissue hydration (22). By eliminating individual degenerative mechanisms, we then created ‘virtual knock-out’ scenarios and investigated their effects on ECM degeneration (15). This provided unique insights into the role of degenerative several mechanisms that drive cartilage degeneration in primary and secondary OA. Still, the current models only account for the degenerative processes that drive cartilage degeneration and overlook the fact that regenerative ECM adaptations also occur under mechanical loading.

In this study, we related physiological mechanical loading to depth-dependent mechanobiological changes in intact and damaged human cartilage explants. We quantified how individual mechanical factors (more specifically maximum shear strain, fluid velocity and dissipated energy) underlie biological adaptations (in this case depth-dependent changes in GAG content) upon mechanical loading. Based on experimental observations of changes in fibril orientation and GAG content upon mechanical loading in cartilage explants, we developed an innovative microstructure-informed FE-based mechanoregulatory algorithm to predict the depth-dependent degeneration and regeneration of GAG content throughout the cartilage thickness upon longitudinal mechanical loading. Following rigorous calibration and validation against longitudinal ex vivo experiments, the model allowed assessment of the impact of compromised tissue integrity on depth-dependent ECM adaptation in the presence of a cartilage defect. As such this predictive mechanobiological model has unique potential to inform and optimize innovative personalized rehabilitation approaches for cartilage preservation.

## Results

### Ex-vivo experimental set-up to evaluate human cartilage adaptations during long term compressive loading

We evaluated the longitudinal biological response of human articular cartilage explants to controlled mechanical stimulation. Therefore, a workflow was established (**Fig. 1A**) to apply long-term, ex-vivo compressive loading to human articular cartilage explants both in the absence and in the presence of a cartilage surface defect.

**Fig. 1.**
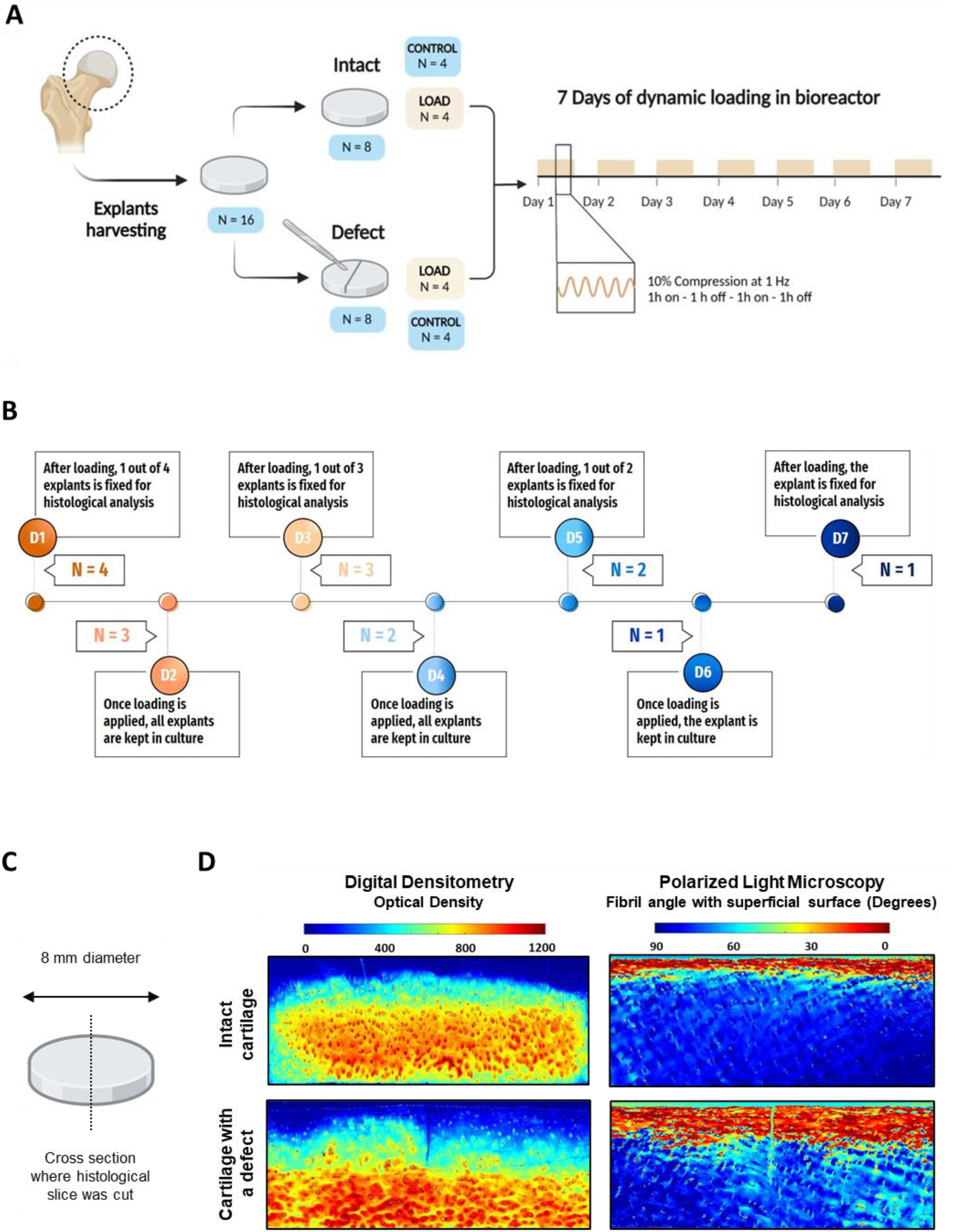
Experimental workflow and example of raw digital densitometry (DD) and polarized light microscopy (PLM) results. (**A**) Schematic of the established workflow for applying long-term, ex-vivo compressive loading to human articular cartilage explants both in the absence and in the presence of a defect. (**B**) Both intact and damaged cartilage explants were loaded for 7 days and fixed for quantitative histological measurements at days 1, 3, 5 and 7. (**C**) Illustration showing a cartilage explant and the location where histological sections were taken, (**D**) Digital densitometry and polarized light microscopy were used to quantify local GAG content and fibril orientation, respectively.

A total of 16 cartilage explants of 8 mm diameter and 2.68 ± 0.46 mm in height were obtained from a hip joint of a female donor with no history or radiographic evidence of OA. A half-depth defect was created in 8 of the explants using a scalpel. The two test groups (intact and defect) were then subjected to physiologic mechanical loading for 1 week using a previously established protocol consisting of 2 cycles per day of 10% compressive strain (compression) at 1 Hz, 1 hour on, 1 hour off. Unloaded explants were kept in culture as controls. At various time-points (**Fig. 1B**), cartilage explants were fixed and sectioned (**Fig. 1C**) for quantitative histological measurements, in particular digital densitometry (DD), and polarized light microscopy (PLM) (**Fig. 1D**). Using these measurements, we documented the time-dependent mechano-adaptations in intact and damaged cartilage tissue in terms of GAG content and fibril orientation throughout the cartilage’s depth upon loading.

### Physiologic mechanical loading maintains GAG content in intact cartilage and cartilage with a defect

To test how physiological mechanical loading for 1, 3, 5 or 7 days affects GAG production in human cartilage explants, we measured GAG content quantitatively by DD in Safranin O-stained cartilage samples. A representative image of the depth-dependent quantitative histology for damaged tissue together with the measured GAG content over tissue depth in the region of interest (ROI), defined as a 2 mm region in the center of the cartilage sample, are shown in **Fig. 2A**.

**Fig. 2.**
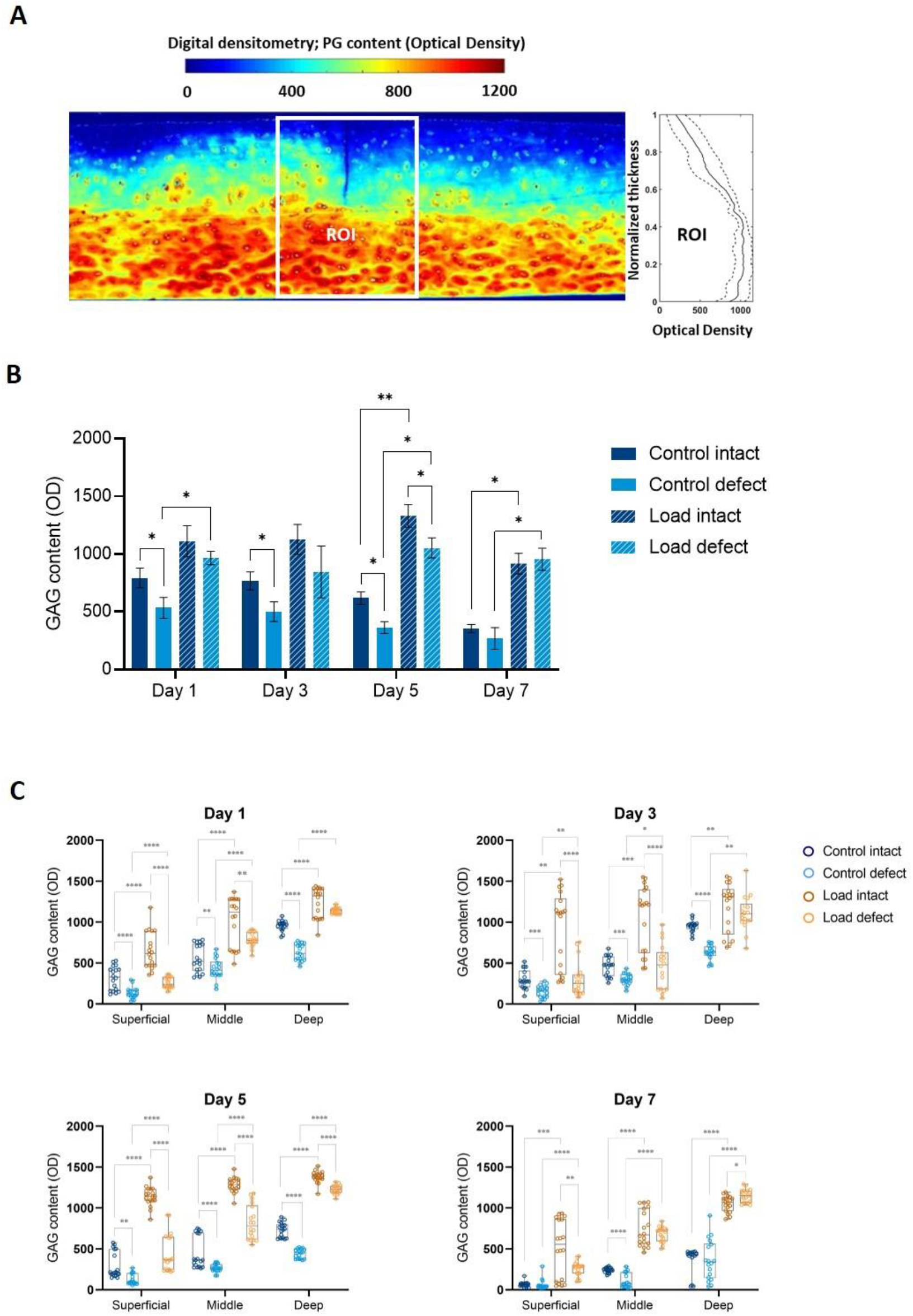
Mechanical stimulation maintains GAG content. **(A)** Representative image of digital densitometry analysis of a cartilage section with a defect and the selected region of interest (ROI). **(B)** Bar graph of GAG quantification in intact cartilage sections and sections with a defect, both in the presence and absence of mechanical stimulation during 7 days of culture (n = 18 measurements per condition). **(C)** Depth-dependent quantification of GAG content in intact cartilage sections and sections with a defect, both in the presence and absence of mechanical stimulation during 7 days of culture (n = 18 measurements per condition). *P* < 0.05 (*), *P* < 0.01 (**), *P* < 0.001 (***), *P* < 0.0001 (****).

In the absence of mechanical stimulation, the average GAG content over the ROI (**Fig. 2B**) decreased over time in both intact explants and explants with a defect. Interestingly, the presence of a defect in control explants significantly decreased the GAG content compared to the intact explants at day 1, day 3 and day 5. Depth-dependent analysis revealed that GAG content in control explants (**Fig. 2C**) increased with cartilage depth, further validating our measurements according to the literature (23, 24).

Upon mechanical stimulation of intact explants, GAG content increased at all time-points and was significantly higher than in the control explants after 7 days of loading and throughout the tissue depth. Surprisingly, mechanical stimulation also increased the GAG content in the samples with a defect, reaching similar levels as intact cartilage explants. However, the presence of a defect resulted in lower GAG content compared to loaded intact explants in all cartilage layers during the first 5 days. Interestingly, the defect-induced loss of GAGs was partially rescued by mechanical loading in all cartilage layers on days 1, 3, 5 and 7.

Taking together, our results suggest that mechanical loading is essential to maintain GAG production to compensate for the defect-induced loss of GAGs and contribute to cartilage homeostasis.

### Presence of chondral superficial defects alters loading-induced fibril reorientation

We investigated the effect of mechanical loading on collagen fibril reorientation. The arcade-like orientation of collagen fibrils plays a crucial role in determining the ability of the tissue to withstand transverse, tensile loads that occur parallel to the cartilage surface upon axial compression. Fibrils oriented parallel to the cartilage surface bear the tensile load, while those oriented more perpendicular to the cartilage surface (high angle) contribute less to tensile strength (25, 26). We evaluated collagen fibril orientation using PLM. A representative image obtained by PLM together with the measured fibril orientation over tissue depth in the ROI are shown in **Fig. 3A**.

**Fig. 3.**
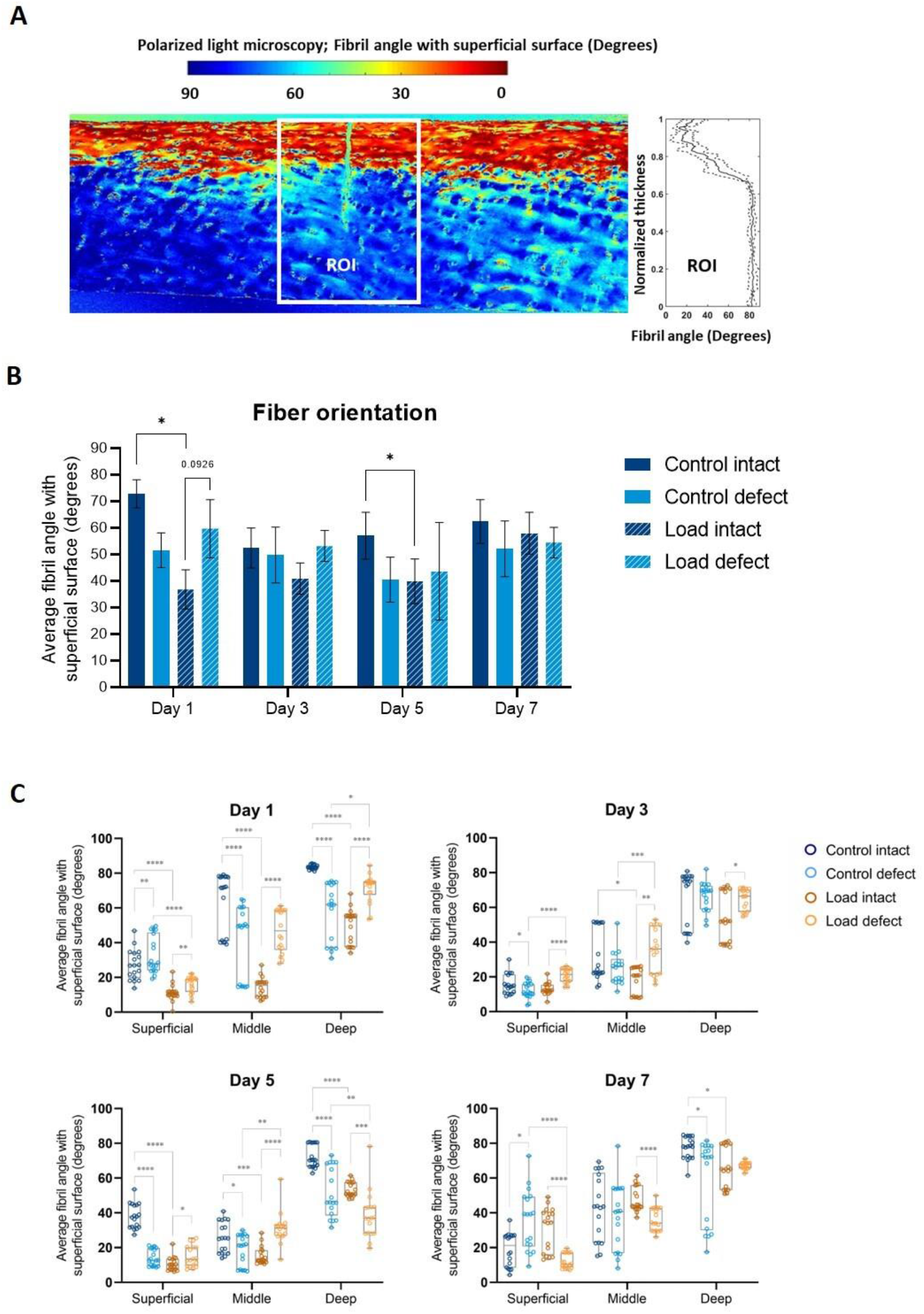
Mechanical loading leads to fibril reorientation. **(A)** Representative image of polarized light microscopy analysis of a cartilage section with a defect and the selected region of interest (ROI). **(B)** Bar graph showing the average fibril angle with the superficial layer in intact cartilage sections and sections with a defect, both in the presence and absence of mechanical stimulation during 7 days of culture (n = 18 measurements per condition). **(C)** Depth-dependent quantification of fibril reorientation in intact cartilage sections and sections with a defect, both in the presence and absence of mechanical stimulation during 7 days of culture (n = 18 measurements per condition). *P* < 0.05 (*), *P* < 0.01 (**), *P* < 0.001 (***), *P* < 0.0001 (****).

In the absence of mechanical stimulation, the presence of a defect resulted in decreased alignment of collagen fibrils with the cartilage surface (**Fig. 3B**), with this effect being significant in middle and deep layers at days 1, superficial and middle layers at day 3, all layers at day 5 and only deep layer at day 7 (**Fig. 3C**).

Mechanical loading induced collagen fibrils to align with the cartilage surface (as reflected by the decreased fibril angle) in intact samples, with this effect being significant at days 1 and 5 (**Fig. 3B**). Depth-dependent analysis shows this effect in all layers up to day 5 (except for the superficial layer on day 3 – **Fig. 3C**). However, disruption of the surface integrity due to a defect impaired the fibril reorientation upon loading as no significant changes in fibril angle were observed, on average in the cartilage explants with a defect. As a result, fibrils were less aligned with the surface in the explants with a defect than intact explants, although not significantly on average (**Fig. 3B**). This suggests that loading in the presence of a defect could lead to disorganization of the collagen network, which may reduce tissue strength.

Taken together, these data indicate that mechanical stimulation induces fibril reorientation in a depth-dependent way in intact cartilage and suggest that the presence of a defect compromises cartilage adaptation by impairing collagen fibril reorientation.

### Mechanoregulatory algorithm relating mechanical micro-environment to load-induced depth-dependent changes in GAG content of intact cartilage

To assess the impact of mechanical micro-environment on GAG content variation, we employed FE models using a fibril-reinforced poroelastic (FRPE) material model that can distinguish the mechanical contribution of cartilage constituents (see Methods section for details). These models were used to estimate dissipated energy (*DE*), maximum shear strain (*MSS*) and fluid velocity (*FV*) – parameters previously associated with anabolic and catabolic responses within the explants (11, 16, 21, 22, 27) – on a daily basis during loading. These models replicated in vitro loading conditions on each loading day. We developed a mechanoregulatory algorithm that relates the outputs of the FE models (*DE, MSS* and *FV*) to the depth-dependent changes in GAG content over a 7-day loading period, aiming to identify the individual contributions of each mechanical parameter.

We calibrated the mechanoregulatory algorithm by optimizing the six weighting factors to match time- and depth-dependent GAG content simulation with the experimental measurements (refer to the Methods section for specifics). Normalizing both the experimental and model-predicted GAG content in loaded samples to that of the corresponding control sample enabled us to accommodate the variability of explants on each loading day and quantify the loading-induced impact on depth-dependent changes in GAG content for each loading day (see **Fig. 4A**).

**Fig. 4.**
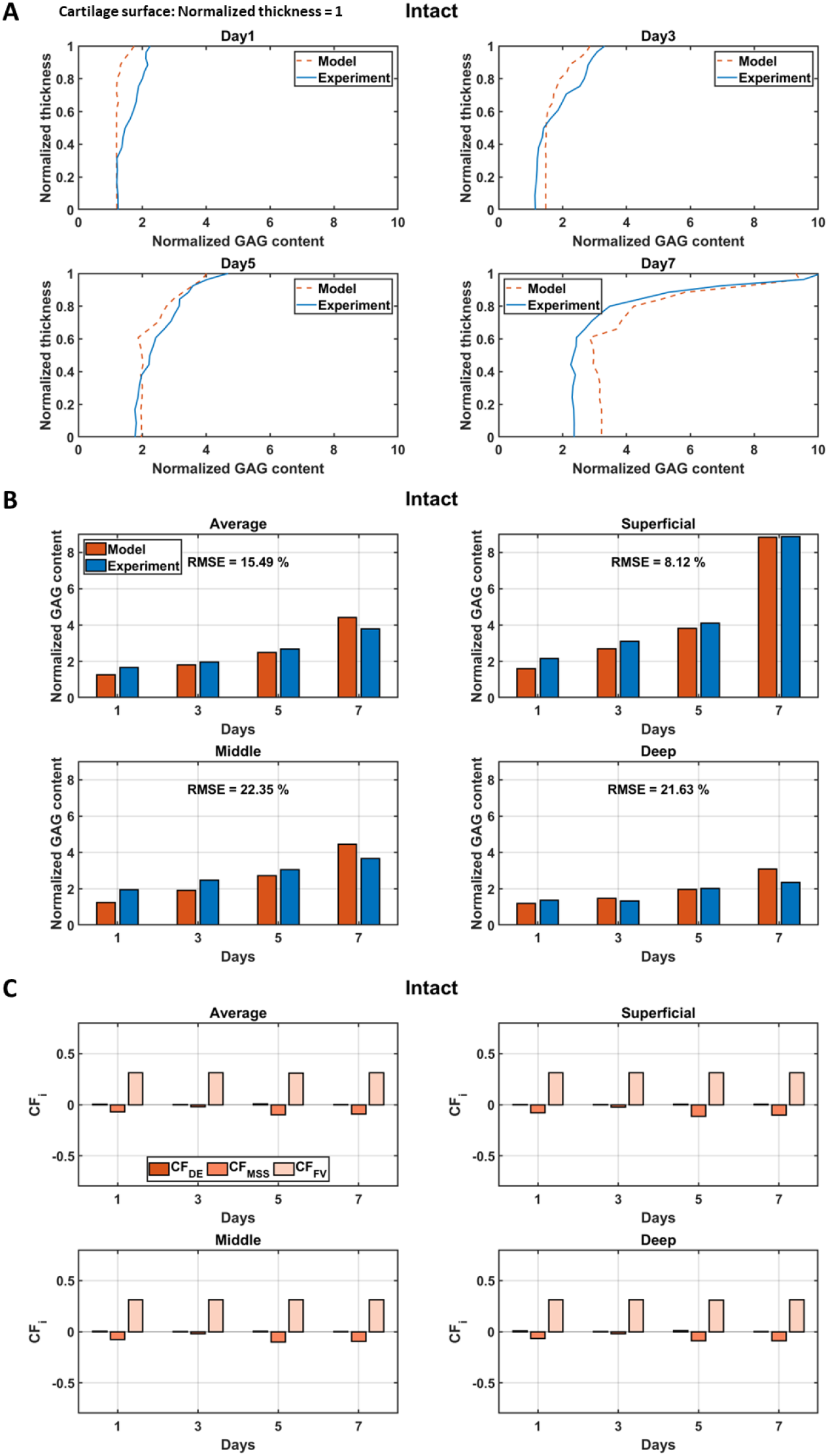
In silico model, calibrated with intact explants, shows the contribution of mechanical micro-environment to the changes in GAG content changes. **(A)** The graph shows the optimal fit of the predicted depth-dependent normalized GAG content by the in silico model over several test days compared to the experimental observations (DD measurements). **(B)** A comparison between the mean value of the predicted normalized GAG content and the corresponding experimental values is shown. This comparison is made over the total cartilage thickness (average) and within the superficial, middle, and deep layers over the course of the test days. The quality of the fit is indicated by the RMSE values. **(C)** Contribution of the three mechanical parameters (FE model outputs: *DE, MSS* and *FV*) to the change in GAG content as reflected in the change factors (*CF*_*i*_ = *CF*_*DE*_, *CF*_*MSS*_ or *CF*_*FV*_).

The calibrated model effectively accounts for depth- and time-dependent changes in GAG content over 7 days of mechanical loading in the intact samples. Specifically, the model exhibits a robust fit with root mean square error of 15.49% for the average GAG content throughout the cartilage thickness (see **Fig. 4B**). Table I shows the values of the identified weighting factors during the calibration process. Notably, the depth-dependent rate of GAG change 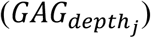 increases from the deep to the superficial zone. More precisely, the rate of GAG change in the superficial zone is approximately twice that in the deep zone, indicating a pronounced depth-dependent variation in GAG turnover within the cartilage tissue. This suggests that mechanically induced changes in GAG content are more prominent in the superficial layer and gradually decrease with cartilage depth.

**Table I:** Calibrated weighting factors and depth-dependent rates of GAG change in the in silico model. These parameters were adjusted to obtain the optimal fit between model predicted and experimental GAG content in intact samples – see eq. (7) and eq. (8).

| Parameter | Identified value following calibration using intact explant DD data |
| --- | --- |
| $k_{DE}$ | 0.63 |
| $k_{MSS}$ | 2.83 |
| $k_{FV}$ | 90.48 |
| $GAG_{depth_{sup}}$ | 1.97 |
| $GAG_{depth_{mid}}$ | 1.38 |
| $GAG_{depth_{deep}}$ | 0.98 |

The parameters within the calibrated mechanoregulatory model enable us to decipher the distinct contributions of individual mechanical triggers to changes in GAG content. In **Fig. 4C** we delineate the individual contribution of each FE model output to the depth-dependent GAG content changes over time, based on their corresponding change factors (i.e. *CF*_*DE*_, *CF*_*MSS*_ and *CF*_*FV*_ that are functions of *DE, MSS* and *FV* defined in eq. (7)). The results reveal that fluid velocity exerts a significant positive influence on GAG production, maintaining a consistent effect across various depths and time points, while dissipated energy exhibits a limited positive effect on GAG production. In contrast, the maximum shear strain negatively affects GAG content.

### In silico mechanoregulatory algorithm can predict depth-dependent changes in GAG content upon loading in cartilage with defect (model validation)

To validate the in silico model predictions, the mechanoregulatory algorithm was used to predict the depth-dependent changes in GAG content over time in defect samples with a surface cut. For this purpose, the weighting factors and depth-dependent rates of GAG change, calibrated for the intact explants (Table I), were used together with the FE models’ outputs (*DE, MSS* and *FV*) of the defect samples. Comparable to the intact samples, the depth-dependent experimental and model predicted GAG content were normalized to the GAG content of the corresponding control samples for each day of loading (**Fig. 5A**). The RMSE of 13.50% for the average of the predicted GAG content over cartilage thickness (**Fig. 5B**) is in the same range as RSME after parameter calibration in the intact samples (15.49% in **Fig. 4B**). This result shows that the in silico model, even when calibrated with the data from the intact samples, is able to predict the mechanics-driven changes induced by a compromised mechanical environment in the defect samples.

**Fig. 5.**
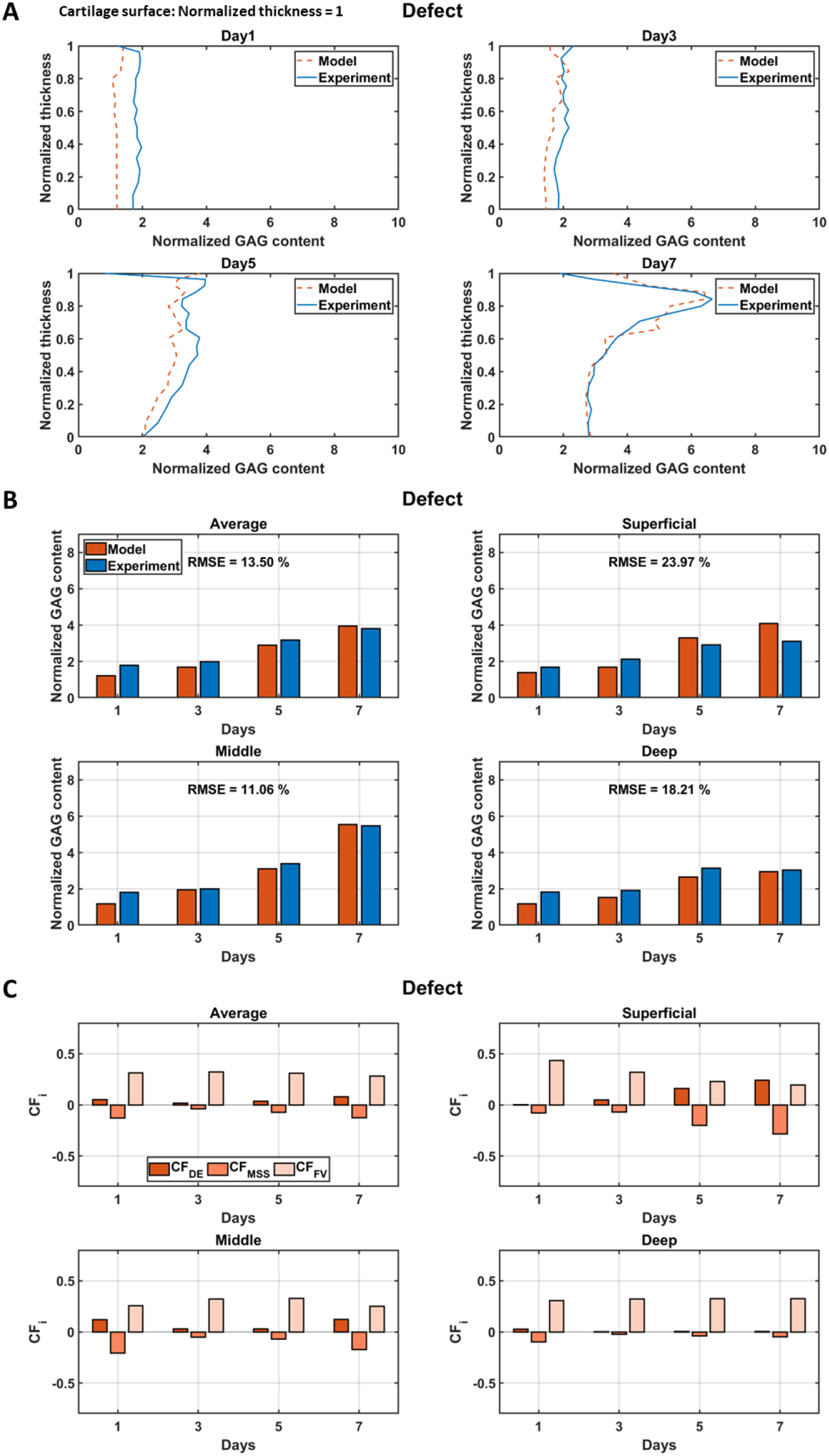
In silico model, validated against explants with defect, can explain and predict depth-dependent changes in GAG content upon loading. **(A)** The graph shows the predicted depth-dependent normalized GAG content by the in silico model over several test days compared to experimental observations obtained through DD measurements. **(B)** A comparison is shown between the mean values of the predicted normalized GAG content and the corresponding experimental values. This comparison is made over the total cartilage thickness (average) and within the superficial, middle, and deep layers over the course of testing days. The RMSEs of the in silico model predictions are shown. **(C)** Contribution of the three mechanical parameters (FE model outputs: *DE, MSS* and *FV*) to the change in GAG content reflected in the change factors (*CF*_*i*_ = *CF*_*DE*_, *CF*_*MSS*_ or *CF*_*FV*_).

To assess whether the compromised mechanical environment in the defect samples affected the contribution of the different FE model outputs to the observed changes in GAG content, we compared the values of their corresponding depth- and time-dependent change factors (i.e. *CF*_*DE*_, *CF*_*MSS*_ and *CF*_*FV*_ that are functions of *DE, MSS* and *FV* defined with eq. (7)) over the measurement days and cartilage thickness (**Fig. 5C**). The results show that, in contrast to the intact samples, *DE* had a significant positive effect on the defect samples in both the superficial and middle zones, with an increasing contribution in the superficial zone over the measurement days. However, the effect was minimal in the deep zone. On the other hand, *MSS* had a negative effect, with its significance decreasing from the superficial to the deep zone and an increasing contribution in the superficial zone over the measurement days. *FV* showed a significant positive effect over the entire depth of the cartilage, but in contrast to *DE* and *MSS*, its effect decreased in the superficial zone over the days of measurement.

## Discussion

Mechanical factors are vital for preserving the structural integrity and functionality of articular cartilage. However, excessive loading or abnormal joint mechanics can hasten cartilage degradation, leading to the development of OA (27). It is imperative to understand which mechanical factors contribute to the degeneration and regeneration of cartilage constituents in health and disease to inform the design of personalized rehabilitation strategies for OA prevention or mitigation.

In this study, we achieved a significant milestone by developing an innovative in silico model that predicts concurrent regenerative and degenerative biological responses of cartilage under longitudinal mechanical loading. Through the identification of mechanical parameters steering these biological changes, we pinpointed the contribution of a novel in silico mechanoregulatory algorithm. The algorithm served as a tool to pinpoint the roles of maximum shear strain, fluid velocity, and dissipated energy as pivotal mechanical parameters influencing depth-dependent ECM adaptation during mechanical loading in an in vitro system.

To this end, in vitro experiments, conducted on intact and damaged human cartilage explants in our bioreactor setup were utilized to respectively calibrate and validate the developed mechanoregulatory algorithm. Calibrated using the intact explants data, the model successfully predicted the overall experimentally observed increase in GAG production within defect explants subjected to mechanical loading (**Fig. 4B** and **Fig. 5B**). Indeed, our experimental results emphasized the necessity of mechanical stimulation to maintain homeostatic GAG levels, not only in intact cartilage but also in cartilage with a defect. Previous evidence supporting the importance of mechanical stimulation for increasing GAG content was therefore consolidated. Our combined approach, blending in vitro experiments with in silico modeling, revealed that physiological cyclic loading optimally increases dissipated energy and fluid velocity, fostering proteoglycan synthesis.

The model demonstrated the ability to predict the depth-dependent adaptation in GAG content of defect cartilage explants following mechanical loading. By incorporating sample-specific, experimentally measured depth-dependent fibril orientation into the finite element models, the depth-dependent rates of GAG change were identified (Table I). The model parameters emphasized the more significant role of mechanics in driving changes in GAG content in the superficial zone compared to the middle and deep zones.

Furthermore, the model showcased its predictive capacity by accurately mirroring the experimentally observed temporal variations in GAG content resulting from mechanical loading of explants with a defect across the cartilage thickness. More specifically, the model can mimic the experimental observation that mechanical stimulation positively influenced GAG content even in the presence of a half-depth cartilage defect, with a more pronounced increase compared to intact cartilage.

The presence of a defect led to distinct GAG content alterations, with less pronounced increases in the superficial layer and more pronounced increases in the middle layer compared to intact explants (solid lines in **Fig. 6**) indicative of a shift in the mechanical environment favorable for GAG production in the middle layer. Remarkably, the in silico model accurately predicted these variations (dashed lines in **Fig. 6**), showcasing its ability to gauge distinct mechanical conditions between intact and defect explants underlying the observed distinct GAG production in the different zones.

**Fig. 6.**
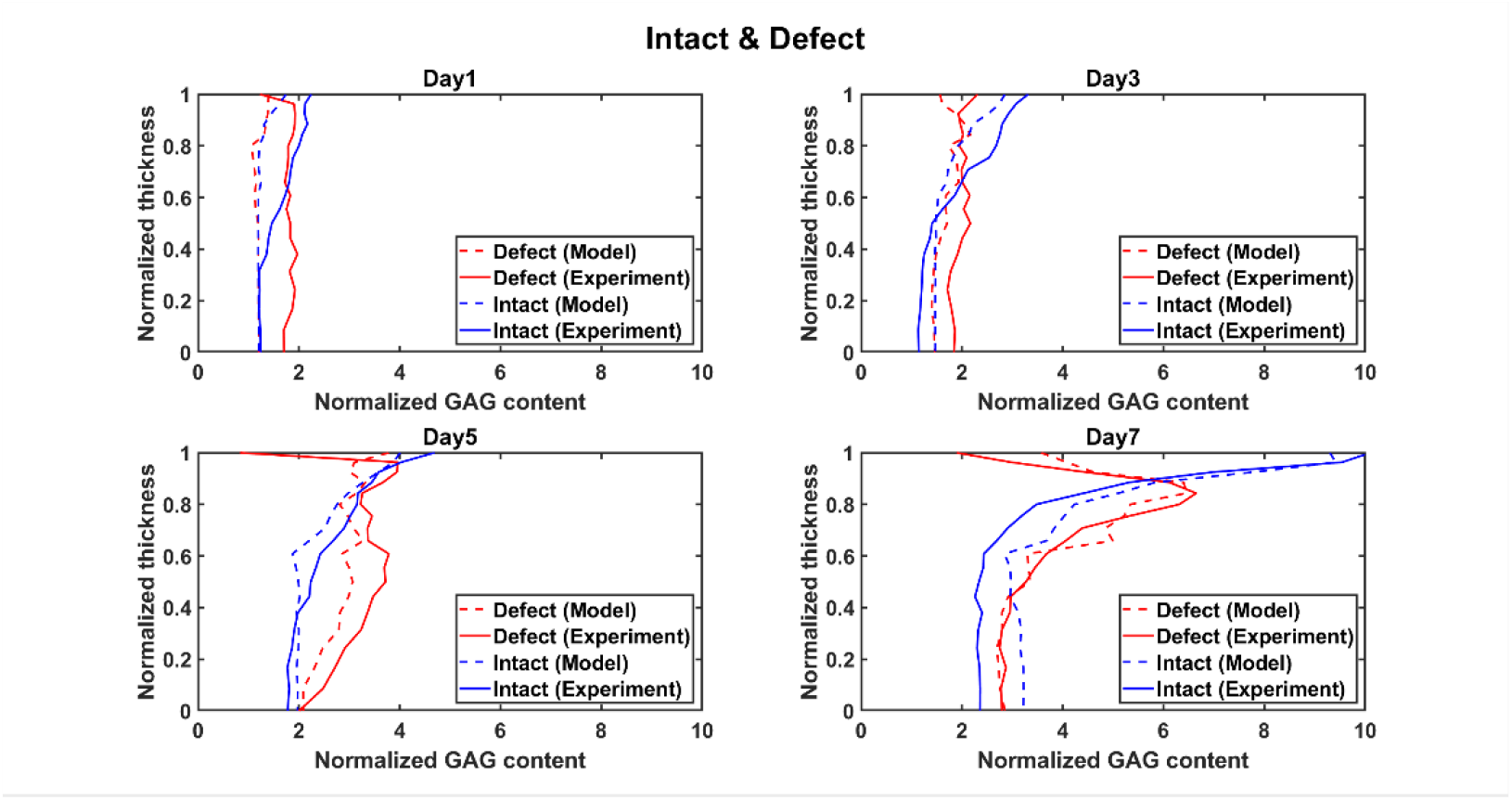
In silico model can predict the temporal variations in depth-dependent GAG content resulting from the mechanical loading and defect. The graph shows the in silico model’s predicted depth-dependent normalized GAG content over multiple testing days compared to experimental observations obtained through DD measurements for intact and defect explants.

Moreover, the mechanoregulatory model showed the unique ability to differentiate the distinct contributions of individual mechanical triggers to GAG content adaptations, which is challenging through current experimental procedures. Comparative analysis revealed *FV* consistently emerging as the main contributor to changes in GAG content induced by physiologic mechanical loading, fostering an increase in GAG production across all layers of intact explants and the deep layer of defect explants (**Fig. 4C** and **Fig. 5C**). Upon closer examination of depth-dependent mechanical contributions, significant differences emerged. In defect explants, unlike intact ones, there was an enhanced regenerative impact of *DE* in the superficial layer (on days 5 and 7) and an enhanced degenerative effect of *MSS* in the superficial layer (on days 5 and 7) and the middle layer (on days 1 and 7). The heightened *MSS*-induced degeneration in the superficial and middle layers of defect explants correlated with increased shear strains due to the defect. These strains, leading to elevated stress, concurrently magnified *DE*’s contribution to GAG production.

To enhance the precision of predictions, a meticulous examination of the calibration process was conducted, assessing fitting quality, prediction errors, and potential discrepancies. The algorithm exhibited exceptional fit in the superficial layer, gradually diminishing with increasing cartilage depth. Although taking into account the sample-specific depth-dependent fibril orientation in the FE models, the predictions demonstrated varying precision across different depths, highlighting the importance of addressing sample-specific depth-dependent fluid content in future work.

To enhance the non-destructiveness and comprehensiveness of our mechanoregulatory model, we propose incorporating cutting-edge experiments using non-destructive imaging techniques and integrating biochemical-driven in silico models. Advanced imaging modalities such as MRI-based methods (28–30) and synchrotron-based imaging (31, 32) promise to refine the model, allowing for more accurate parameter calibration. These techniques can offer valuable insights into cartilage microstructure strain and 3D composition under mechanical loading. Their non-destructive nature allows for longitudinal tracking of changes in cartilage properties within a single sample. The model’s current limitation in considering the impact of biochemical factors observed in OA joints, needs to be acknowledged suggesting integration with existing in silico models focusing on biochemical and inflammatory aspects (33–35) in order to evolve to a comprehensive mechano-biochemical model.

Given our focus on developing the FE-based mechanoregulatory algorithm and its calibration and validation, along with the constraints of accessing non-OA human cartilage samples and conducting time-intensive and expensive longitudinal mechanical loading experiments, we used cartilage samples from a single donor limiting the generalizability of the experimental results. Therefore, for robust and conclusive statistical analysis of the experimental findings, future studies should consider increasing the sample size. This approach would enable incorporating probabilistic modeling approaches to address input parameter variability, hence leveraging probabilistic instead of deterministic model outcomes.

Ultimately, the in silico mechanoregulatory model holds significant potential to impact in vitro and in vivo experiments, unraveling the intricate interactions between mechanical loading and biological changes. In the context of tissue engineering, the model can optimize interventions by identifying the role of individual mechanical parameters in ECM adaptations under mechanical loading. This not only enhances therapy effectiveness but also contributes to reducing reliance on animal experiments, addressing ethical concerns associated with them. Additionally, the model offers opportunities to optimize in vitro mechanobiological experiments, aiding in designing new OA rehabilitation approaches. This aligns with recent endeavors to advance in silico trials, which aim at simulating clinical conditions, predicting treatment outcomes, and thereby facilitating more informed and efficient clinical trials. As such, this work underscores the future potential of the model in understanding clinical conditions, predicting outcomes and advancing personalized rehabilitation and medicine approaches.

## Materials and Methods

### Experiments

#### Human articular cartilage harvesting and creation of defect

Informed consent and ethical approval of the University Hospitals Leuven Ethics Committee and Biobank Committee (Leuven, Belgium) (S56271) was obtained before performing this study. A total of 16 healthy human cartilage explants were harvested from the hip of an 80-year-old female donor with no history of OA, undergoing hip replacement surgery for osteoporotic or malignancy-associated fracture. First, cartilage was dissected into fragments using a scalpel and then rinsed in phosphate buffered saline (PBS) supplemented with 1% (vol/vol) antibiotic/antimycotic (Gibco). The cartilage fragments were subsequently cut with an 8 mm diameter disposable biopsy punch (Robbins Instruments). A cut (defect) through half of the depth of the explant was performed with a scalpel and using a metal ring to keep constant the depth of the defect among explants. All samples were cultured for 24h in DMEM/F12 (Gibco), 10% foetal bovine serum (FBS, Gibco), 1% (vol/vol) antibiotic/antimycotic (Gibco), and 1% L-glutamine (Gibco) at 37°C and 5% CO2 humidified atmosphere before starting the loading experiments.

#### Dynamic loading in bioreactor

Each day before starting the loading, the cartilage explants’ thickness was measured in a Leica M50 Stereo Microscope equipped with a DFC425 C Digital Microscope Camera. Ex vivo mechanical stimulation was applied to articular cartilage explants in the ElectroForce® BioDynamic 5210 bioreactor (TA Instruments) with the accompanying WinTest® Software, inside an incubator at 37°C and with 5% of CO2. All elements of the loading chambers were gas sterilized before use. Cartilage explants were positioned in the loading chamber with the upper layer of the cartilage facing up, while keeping sterile conditions. The explant was secured between the two surfaces of the sample holder. Once closed, the loading chambers were filled with culture medium (DMEM/F12 + 10% FBS + 1% antibiotic/antimycotic + 1% L-glutamine) and placed in the bioreactor. The applied loading protocol consisted of 1 hour of 10% sinusoidal compressive strain (compression) at 1 Hz + 1 hour of free swelling + 1 hour of 10% compression at 1 Hz + 1 hour of free swelling. 10% compressive strain was defined based on the measured sample thickness. For each donor, paired explants were maintained under free swelling conditions in an incubator at 37°C and 5% CO2 as unloaded controls.

#### Quantitative measurements of GAG content and collagen fibril orientation

On day 1, day 3, day 5 and day 7, two loaded explants (one intact and one with a defect) and two unloaded controls (one intact and one with a defect) were harvested for quantitative histological measurements. Explants were rinsed with PBS and fixed with 4% formaldehyde in PBS for 24 h at 4°C. Then, samples were decalcified in 0.5 M EDTA in PBS pH 7.5 for 5 days, changing the EDTA solution every day, at 4°C. On the 6^th^ day, explants were washed in running tap water for 3 hours and submerged in methanol for 24 hours prior to paraffin embedding. Cartilage sections of 3 µm were prepared for staining and 18 sections were prepared per sample. DD and PLM measurements were performed as described previously (36). Briefly, for measurements of GAG content by DD, sections were dewaxed, rinsed in tap water, and stained with safranin-O for 5 min. After a 5 min wash under running tapwater, slides were air dried for 10 min, dehydrated with 4 washes in ethanol and mounted on a microscope slide. Stained sections were then photographed using a standard light microscope (Nikon Microphot FXA, Nikon Co., Tokyo, Japan) with a pixel size of 3.09 × 3.09 µm. Imaging was performed with a CCD camera (Hamamatsu ORCA-ER, Hamamatsu Photonics, Hamamatsu, Japan) and a monochromator (λ = 492 ± 5 nm), which was calibrated using neutral filters spanning a wide range of optical densities from 0 to 3.0.

Unstained sections were used for the evaluation of fibril orientation angle by polarized light microscopy (PLM). The custom-designed PLM system consisted of a microscope body (Leitz Ortholux II POL light microscope, Leitz Wetzlar, Wetzlar, Germany), light source with a monochromator (λ = 630 ± 30 nm, Edmund Optics Inc., Barrington, NJ, USA), crossed polarizers (Techspec optics® XP42-200, Edmund Optics, Barrington, NJ, USA) and a monochrome camera (BFS-U3-88S6M-C FLIR Blackfly® S, FLIR Systems Inc., Wilsonville, OR, USA, pixel size 3.5 µm × 3.5 μm) with a 2.5× magnification lens. The collagen fibrils and the collagen network structure induce an angle dependency on the observed light intensity. This allows the determination of pixel-wise fibril angle and parallelism index (anisotropy) via Stokes parameters and Michelson’s contrast method (37).

For each sample, we examined the measured GAG content and fibril orientation data using a rectangular 2 mm wide ROI spanning throughout the tissue thickness (**Fig. 2A** and **Fig. 3A**): the ROI was positioned in the middle of explant for the intact explants, and it was positioned symmetrically to the cut for the defect explants. Depth-wise profiles were then obtained by averaging over 18 sections within the ROI (Matlab R2022b, Mathworks Inc., MA, USA). Additionally, based on the fibril orientation data of the day 1 control sample, the normalized thicknesses of the superficial, middle, and deep zones were 0.10, 0.25 and 0.65, respectively. These thickness values were subsequently used to divide the sections into the three zones, allowing for the averaging of GAG content and fibril orientation within each zone in the ROI for further depth-wise analysis (see **Fig. 2A**).

#### Statistical analysis

Data analysis and graphical presentation were performed with GraphPad Prism version 8. Data are presented as mean ± SEM and as individual data points, as indicated in the figure legends. GAG content and fibril angle data averaged over depth were compared between conditions at each time point (**Fig. 2B** and **Fig. 3B**) or cartilage layer (**Fig. 2C** and **Fig. 3C**) by paired 2-tailed Student’s test or Wilcoxon test in case of non-normal distribution. P values of pairwise comparisons are indicated in the graphs. Model assumptions were further checked by a Shapiro-Wilk test, QQ plots and homoscedasticity plots. The homogeneity of variance was evaluated by standardized residuals versus fit plot. P values of less than 0.05 were considered significant.

### In silico modeling

#### Sample-specific FE models

To assess the specific mechanical parameters that influence GAG content adaptations across the cartilage geometry, FE models were constructed for the eight loaded cartilage explants, consisting of four intact and four defect samples (**Fig. 7A**). The sample-specific models comprised of cylindrical geometries with dimensions determined from measurements in the stereomicroscope. For modeling the defect samples, a surface cut of 20 μm width was created, comparable to the thickness of the scalpel used to create the surface cut on the defect explant, extending until half of the explant model thickness.

**Fig. 7.**
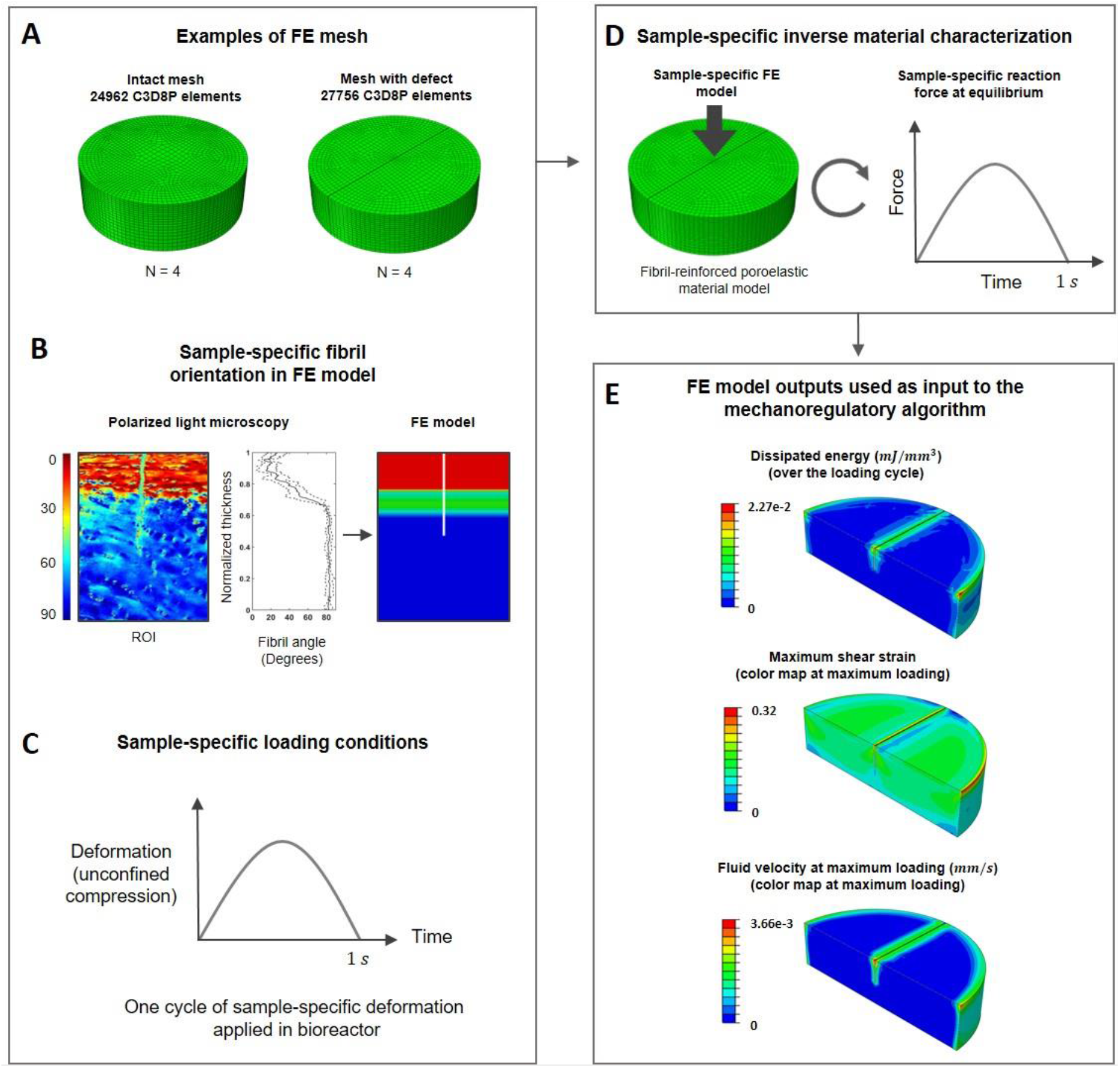
Sample-specific FE modeling workflow. **(A)** Examples of FE mesh, **(B)** sample-specific fibril orientation implemented in the FE model from the PLM measurements, **(C)** one cycle of sample-specific unconfined compression deformation applied to the explant, **(D)** Inverse material characterization to obtain the sample-specific material constants of FRPE material model and **(E)** examples of the three FE model outputs over the cartilage explants.

To model the mechanical behavior of articular cartilage, a 3D FRPE material model was used (38). The FE models were implemented with sample-specific depth-dependent fibril orientation using data from PLM measurements (**Fig. 7B**). Depth-dependent fluid content, decreasing from the top surface to the bottom surface, was based on the literature (22), as we did not have the sample-specific data. The bottom surface of the explants was constrained against vertical translation, while both top and bottom surfaces were permitted to undergo radial expansion. A single cycle of sample-specific sinusoidal compressive deformation, in accordance with the applied deformation within the bioreactor, was applied to the top surface of the model (**Fig. 7C**). This loading was used as a representative of 1 day of the in vitro loading experiment. The fluid was allowed to flow through the side surface (pore pressure = 0).

The intact and defect samples were meshed using 24962 and 27756 linear pore pressure continuum elements (element type C3D8P), respectively (Abaqus/Standard software 2022 by Dassault Systèmes Simulia Corp, USA – **Fig. 7A**). A biased mesh seeding was employed to achieve finer element sizes near the open surfaces of the model, where fluid pressure and deformation gradients are larger. A mesh convergences analysis was conducted, using half, twice, and four times the selected number of elements. Simulations with higher mesh densities revealed no significant differences in deformation gradient, stress and fluid velocity distributions, indicative that the FE model outcomes (i.e. *DE, MSS*, and *FV*) used within the mechano-regulatory algorithm were not affected.

To identify sample-specific material properties – i.e. the constants of the FRPE material model (38) – an inverse parameter fitting approach was employed (**Fig. 7D**). The reaction force calculated at the bottom surface of the FE models was iteratively matched to the measured reaction force of the corresponding cartilage sample within the bioreactor by altering the material constants (details in (38)). For the parameter fitting, the measured force after reaching equilibrium during the second loading of each sample on the day of harvesting for histological measurements was used. To this end, the “lsqnonlin” optimization function (Matlab software R2022b by Mathworks Inc., MA, USA) was used.

Three mechanical parameters, namely the *DE, MSS*, and *FV*, were recorded as outputs from the FE models across the explants’ geometry (**Fig. 7E**) to be later associated with GAG content measurements. Both the *MSS* and *FV* were recorded at each integration point and time increment throughout the loading cycle, with specific information about the calculation method for *MSS* available in (22). The *DE* per unit volume was calculated over the loading cycle by summing the absolute areas of hysteresis loops obtained for the components of stress versus deformation gradient in each integration point. The three parameters were averaged over the cartilage thickness within the specific sample region (ROI as shown in **Fig. 2A**).

#### Mechano-regulatory algorithm

First, a non-localization theory as described in (22, 39) was used to avoid localization of FE model outputs. The non-localized FE model output at each intended integration point (*ip*) of the FE model was obtained as:

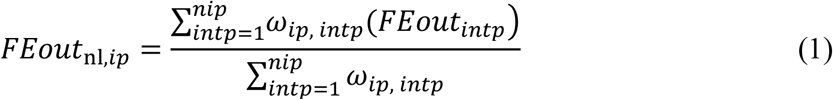

where, *FEout*_nl,*ip*_ is the non-localized FE model output that can be *DE*_nl,*ip*_, *MSS*_nl,*ip*_ or *FV*_nl,*ip*_ and *intp* and *nip* are the index and the total number of integration points in the FE model. ω_*ip*, *intp*_ is the Gauss weighting function at the intended integration point (*ip*) with respect to other integration points (*intp*) and was obtained as:

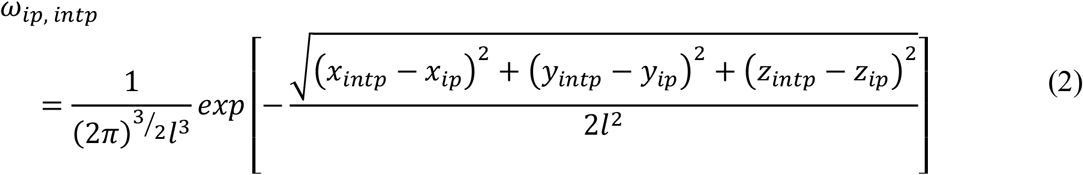

where *x*_*j*_, *y*_*j*_, and *z*_*j*_ are the coordinates of intended and other integration points and *l* is the characteristic length, which is a property related to the scale of the microstructure. This parameter was selected to be equal to the superficial layer thickness (39).

The *DE*_nl,*ip*_ was obtained over the load-unload cycle while *MSS*_nl,*ip*_ and *FV*_nl,*ip*_ were averaged over loading duration. Therefore, the time average of *MSS*_nl,*ip*_ and *FV*_nl,*ip*_ in the load-unload cycle were calculated as:

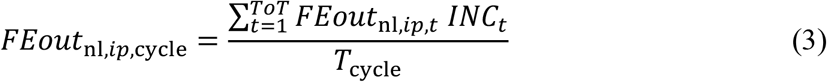

where *FEout*_nl,*ip,cycle*_ can be *MSS*_nl,*ip,cycle*_ or *FV*_nl,*ip,cycle*_. *T*_*cycle*_ is the total time of the loading cycle. *TOT* and *INC*_*t*_ are the total number of time points (*t*) during each loading step and the duration of each time increment in the FE model. *FEout*_nl,*ip,t*_ is the FE model output in each time increment of the FE model, which can be *MSS*_nl,*ip,t*_ or *FV*_nl,*ip,t*_.

Subsequently, to ensure comparability with the histological measurements, the depth-dependent, non-localized FE model outputs were obtained. This was achieved by averaging their values over the same region of interest (ROI) as defined for the histological sections (refer to **Fig. 2A** and **Fig. 3A**). Drawing inspiration from a mechano-adaptive model proposed for tendon tissue (*40*), the resulting averaged parameters were employed to establish three normalized factors (as provided in eq. (4) to eq. (6) and visualized in **Fig. 8**), indicative of their positive or negative effects on the GAG content.

**Fig. 8.**
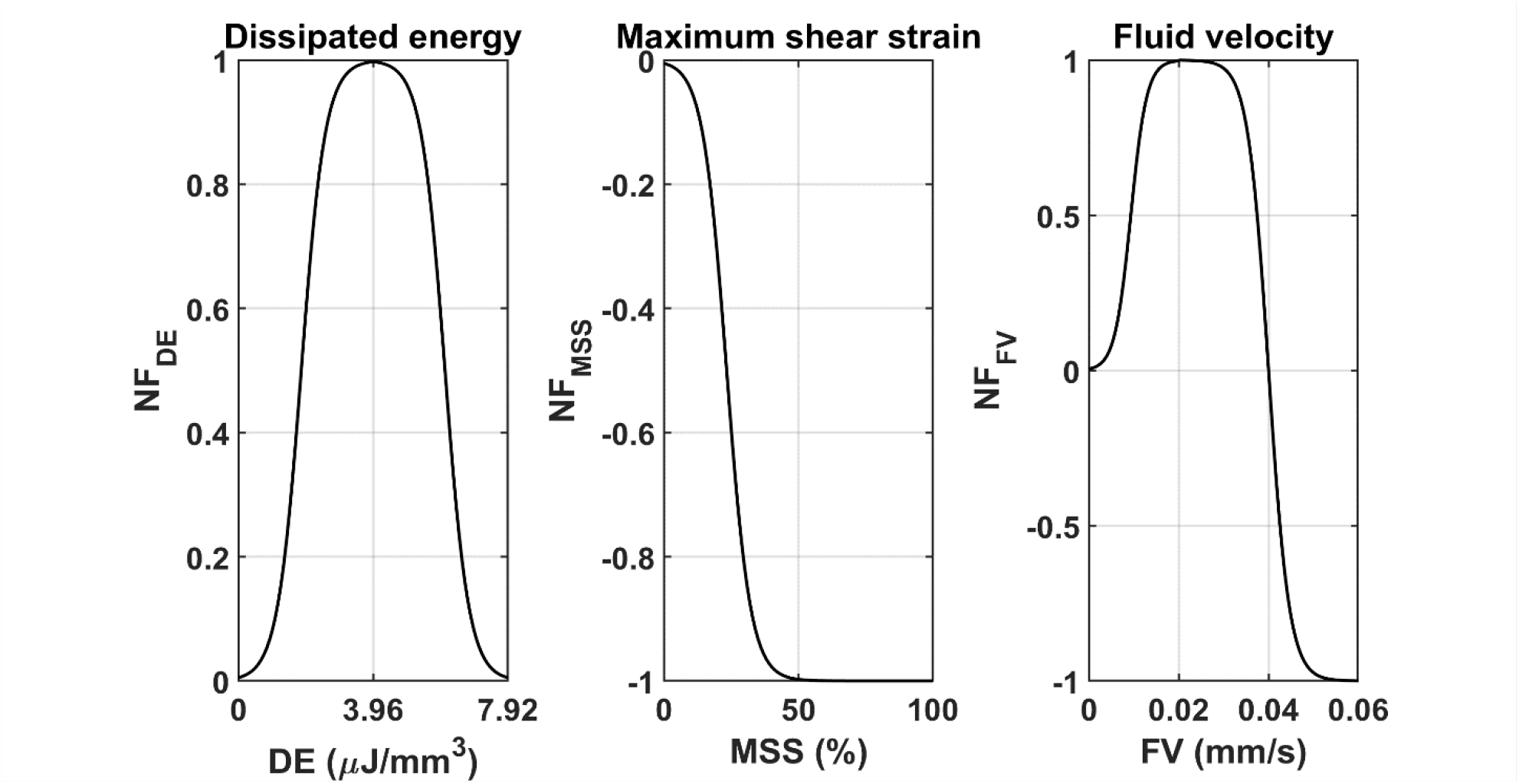
Normalized effects of FE model outputs on the change in GAG content demonstrated by normalized factors. The curves were obtained by plotting the *NF*_*DE*_, *NF*_*MSS*_ and *NF*_*FV*_ versus *DE, MSS* and *FV* using eq. (4) to eq. (6). Positive and negative values demonstrate GAG production and degeneration effects respectively.

As suggested in the literature (11, 12, 41), dissipated energy can stimulate the cells for further GAG production. Therefore, its positive effect was simulated as:

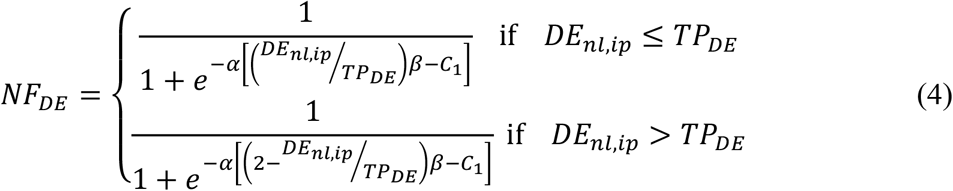

where, *α =* 75, *β =* 0.15, *C*_1_ *=* 0.07 are shape parameters (40) and 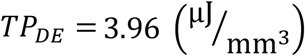 is a transition point at which the dissipated energy stimulates the maximum GAG production. This value was determined by averaging the dissipated energy over a loading cycle during the equilibrium state of intact cartilage explants on the initial day of loading within the bioreactor. This condition is regarded as the closest approximation to the mechanical behavior of native healthy cartilage.

According to the literature (21, 22, 42), maximum shear strain is associated with PG depletion and loss of GAG chains. Therefore, we defined it’s negative effect on GAG content as:

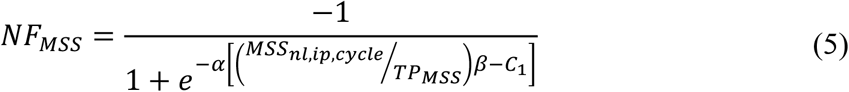

where, *TP*_*MSS*_ *=* 0.5 represents the point at which *MSS* reaches its most significant negative impact (maximum degeneration) on GAG content, a value obtained from a preceding study (21).

Fluid flow within the tissue can yield either positive or negative outcomes on GAG content. A moderate fluid flow (moderate *FV*) has the potential to facilitate nutrient and growth factors delivery while imposing a moderate hydrostatic fluid pressure on cells, believed to enhance chondrocyte GAG production activity (10). Conversely, excessive fluid velocity can harm cells and elicit the extraction of small GAG protein fragments (*21*). We integrated the impacts of *FV* on GAG content as follows:

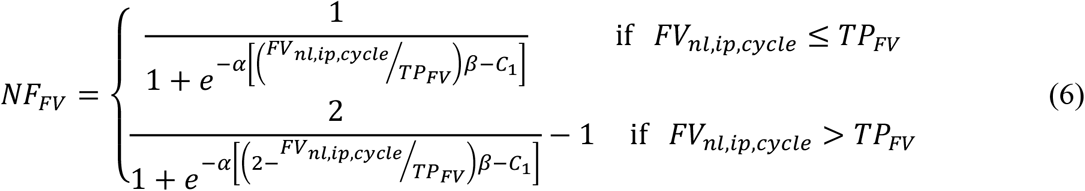

where, *TP*_*FV*_ *=* 0.02 (mm/s) represents the transition point at which *FV* has its maximum positive effect on GAG production and assumed as half of the degradation threshold (*FV =* 0.04 (mm/s)) proposed in an earlier study (21).

To calculate the effect of each FE output on the change in GAG content, the normalized factors were multiplied to weighting factors and change factors were obtained as:

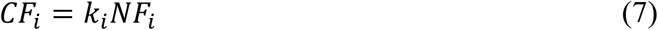

where, *k*_*i*_ represents the weighting factors of the FE model outputs (i.e. *k*_*DE*_, *k*_*MSS*_ and *k*_*FV*_). Therefore, *CF*_*i*_ represents three distinct change factors related to each of the FE model outputs (i.e. *CF*_*DE*_, *CF*_*MSS*_ and *CF*_*FV*_). These factors indicate the significance of each FE model output in the overall change in GAG content due to mechanical loading. To reflect the experimentally observed depth-dependent changes in GAG content, the change factors were summed and multiplied to a depth-dependent rate of GAG change and the total change factor was obtained as:

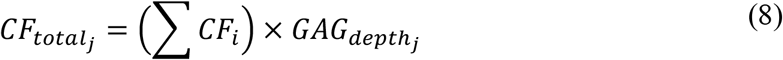

where, 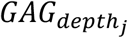 represents the depth-dependent rate of change in GAG content where *j* can be the superficial, middle or deep layer (i.e. 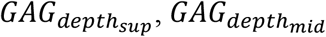 and 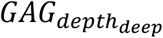).

Finally, the depth-dependent GAG content in each day of loading was predicted as:

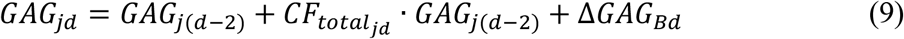

where, *d* represents the day of loading in the bioreactor (days 1, 3, 5 and 7) and *ΔGAG*_*Bd*_ is the basal change in the GAG content. Given that the present mechano-regulatory model primarily aims to replicate the GAG content variations driven by mechanical parameters, *ΔGAG*_*Bd*_ encompasses the free swelling effect on GAG content within the explants. This parameter was derived by calculating the difference between the depth-dependent GAG contents of the control samples across two consecutive DD measurement days. The GAG content at day zero (*GAG*_*0*_), before the first loading, was considered as the measured GAG content using DD in the control sample at day 1.

#### Calibration, validation and sensitivity analysis of mechano-regulatory algorithm

To calibrate the mechano-regulatory algorithm, the six unknown parameters within eq. (7) and eq. (8) – namely *k*_*DE*_, *k*_*MSS*_, *k*_*FV*_, 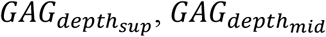, and 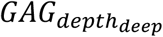 – were identified by fitting these parameters to optimally match the GAG content predictions from the mechano-regulatory algorithm to the experimental DD measurement results of the intact explants. To this end, the obtained outputs from the FE models of intact explants were fed into the mechano-regulatory algorithm, initially employing an estimated set of the six parameters. To account for the variability in the explants at each loading day and estimate the loading effect on the changes in GAG content, both the experimental and the predicted GAG content by the model in the loaded samples were first normalized to the GAG content of the corresponding control sample and the normalized values were compared. Through an iterative process using the “lsqnonlin” optimization function in Matlab (Mathworks), these six parameters were optimized to fit the normalized GAG contents obtained from the algorithm to the corresponding experimental values. During the fitting process, we considered the depth-dependent effect by simultaneously fitting the GAG contents over the cartilage thickness within the ROI (see **Fig. 2A**).

To validate the mechano-regulatory algorithm, the identified parameters were applied in conjunction with FE models of the defect explants to predict GAG content in the loaded samples across the loading days in case of an impaired mechanical environment. These results were then compared to the corresponding DD measurements.

Finally, the calibrated and validated mechano-regulatory algorithm enabled a parameter sensitivity analysis to explore the impact of each FE model output on mechanics-driven changes in GAG content – *i*.*e. CF*_*i*_ values in eq. (7). This entailed a comparison of average *CF*_*i*_ values over the cartilage thickness as well as their average across the superficial, middle and deep zones on the loading days.

## Acknowledgments

We are grateful to the UZ Leuven nursing staff for their efforts to provide cartilage samples for this work and to Lies Storms for her help with cartilage sample preparation for histology.

## Author contributions

Conceptualization: SAE, RCV, IJ

Methodology: SAE, RCV, PT, LM

Investigation: SAE, RCV

Visualization: SAE, RCV

Supervision: NF, RK, IJ

Writing—original draft: SBB, DLA

Writing—review & editing: SAE, RCV, PT, LM, NF, RK, IJ

## Data and materials availability

The raw data supporting the conclusions of this article will be made available by the authors upon request.

